# Human-Genetic Ancestry Inference and False Positives in Forensic Familial Searching

**DOI:** 10.1101/2020.03.06.981134

**Authors:** Alyssa Lyn Fortier, Jaehee Kim, Noah A. Rosenberg

**Author notes:** To whom communication should be addressed (Noah A. Rosenberg).

## Abstract

In forensic familial search methods, a query DNA profile is tested against a database to determine if the query profile represents a close relative of a database entrant. One challenge for familial search is that the calculations may require specification of allele frequencies for the unknown population from which the query profile has originated. Allele-frequency misspecification can substantially inflate false-positive rates compared to use of allele frequencies drawn from the same population as the query profile. Here, we use ancestry inference on the query profile to circumvent the high false-positive rates that result from highly misspecified allele frequencies. In particular, we perform ancestry inference on the query profile and make use of allele frequencies based on its inferred genetic ancestry. In a test for sibling matches on profiles that represent unrelated individuals, we demonstrate that false-positive rates for familial search with use of ancestry inference to specify the allele frequencies are similar to those seen when allele frequencies align with the population of origin of a profile. Because ancestry inference is possible to perform on query profiles, the extreme allele-frequency misspecifications that produce the highest false-positive rates can be avoided. We discuss the implications of the results in the context of concerns about the forensic use of familial searching.

## Introduction

In forensic genetics, when no exact match of a DNA profile to an entrant in a database of profiles can be found, investigators can often test for partial matches to determine if a sample of interest might be a close relative of a database entrant (Bieber *et al*., 2006; Gershaw *et al*., 2011; Butler, 2012). If a partial match is identified, then investigators can consider relatives of the match as possible contributors of the query profile.

Much of the discussion surrounding the suitability of this familial search technique in forensic genetics has centered on the problem of false-positive relatedness matches (Greely *et al*., 2006; Murphy, 2010; Rohlfs *et al*., 2012, 2013; Garrison *et al*., 2013). In searches for exact matches, a sample is typically tested at a number of forensic DNA markers that is small, but large enough that a false-positive database match of a non-contributor to the query at all typed loci is relatively unlikely. In familial identification, however, for a fixed set of markers, because a true relative of the contributor of the query profile has only a partial match, the chance of a false positive—the probability that a non-relative also achieves this less stringent partial match threshold—greatly exceeds the probability that the same non-relative is a false exact match. Hence, owing to nontrivial false-positive rates, close relatives of database entrants can be exposed to inappropriate forensic investigation when they have not in fact contributed to query profiles.

Accurate understanding of the magnitude of false-positive rates in familial search is important for discussions regarding appropriate use of the technique. To study properties of the false-positive rate in familial identification, Rohlfs *et al*. (2012) focused on the choice of allele frequencies used as part of familial-search likelihood calculations. Because a query profile represents a sample from an unknown individual, its population membership, and hence, the appropriate choice of allele frequencies for the calculation, is not known and can potentially be misspecified. With a goal of examining the effect of misspecifying the allele frequencies, Rohlfs *et al*. (2012) used allele-frequency data for a variety of populations to measure rates at which false partial matches between pairs of individuals were identified under a sibling relationship hypothesis when the individuals were in fact unrelated. Rohlfs *et al*. (2012) examined false positives under each of several possible misspecifications, finding that false positives were more likely with misspecified frequencies than when the frequencies were properly specified to correspond to the population of origin of the individuals—especially as the magnitude of the misspecification increased to represent genetically distant populations.

We propose that the most extreme allele-frequency misspecifications that produce the highest false positive rates are possible to avoid by use of an ancestry-inference step in the familial search procedure. Forensic genetic profiles, even with the relatively limited marker sets they typically employ, contain considerable information about genetic ancestry (Phillips, 2015; Algee-Hewitt *et al*., 2016). Thus, if the genetic ancestry of a query profile can be partially inferred prior to a familial search, then the allele frequencies used in the search could be selected as those relevant to the estimated ancestry. Provided the estimated ancestry information is reasonably accurate, extreme misspecifications and the high false positive rates that result from them might be avoided.

Here, we devise a scheme that first infers the genetic ancestry of a query profile and then applies the allele frequencies of the inferred population of origin in familial search computations. Applying this scheme to samples from diverse populations, the false positive rates we observe with the ancestry-inference step are substantially lower than those seen by Rohlfs *et al*. (2012) with misspecified allele frequencies. In fact, they are close to the lower false-positive rates seen by Rohlfs *et al*. (2012) in scenarios with allele frequencies associated with the source population for the query profile. Thus, use of ancestry inference can potentially place an upper bound on the false positive rates of familial search procedures. We discuss the findings in relation to ongoing arguments about the utility and application of familial search.

## Materials and Methods

### Data

We examined a sample of 978 individuals from the Human Genome Diversity Panel (HGDP), genotyped at 791 microsatellite (STR) loci: 13 CODIS loci used in forensic genetics and 778 non-CODIS loci. The data are taken from Algee-Hewitt *et al*. (2016), dropping duplicate locus *TPO-D2S* as in Edge *et al*. (2017). We grouped the individuals into four population groups: Sub-Saharan African (A), European, Middle Eastern, and Central/South Asian (EMC), East Asian and Oceanian (EAO), and Native American (NA). These four groups approximate four clusters that are somewhat genetically distinguishable with the 13 CODIS loci (Algee-Hewitt *et al*., 2016). The numbers of individuals genotyped were 94, 532, 269, and 83, for A, EMC, EAO, and NA, respectively.

### Ancestry Estimation

We performed ancestry estimation using STRUCTURE (Pritchard *et al*., 2000), employing unsupervised clustering with the admixture and correlated allele frequencies models. All STRUCTURE runs used *K* = 4 and a burn-in period of 10, 000 steps followed by 10, 000 iterations from which posterior distributions were calculated. We performed STRUCTURE runs separately using the full set of 791 loci and only using the 13 CODIS loci, in each case employing 10 replicate analyses with the same settings. We averaged the resulting estimated ancestry proportions and estimated cluster allele frequencies using CLUMPP (Jakobsson and Rosenberg, 2007) with the greedy algorithm (*M* = 2), greedily aligning runs in each of 10,000 sequences (*GREEDY OPTION* = 2, *REPEATS* = 10000), and employing the *G* statistic (*S* = 1). We used DISTRUCT to visualize the ancestry estimates (Rosenberg, 2004).

### Simulating Relatives

Each STRUCTURE replicate run using all 791 loci provided estimates of the allele frequencies at each locus for each of the four inferred clusters. Taking the CLUMPP average across the 10 replicate runs, we extracted the estimated allele frequencies, 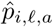, for each cluster *i*, locus *ℓ*, and allelic type *a*. For each of the 978 individuals, to simulate relatives of the individual, we weighted these estimated allele frequencies by the individual’s estimated membership proportions 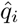, averaged over the 10 replicate STRUCTURE runs with 791 loci, to obtain an appropriate allele frequency distribution for each individual, as in Equation 1:

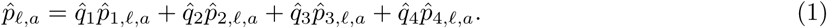

For each of the 978 individuals, we simulated 10 siblings. To generate each sibling, at each locus, we copied both of the original individual’s alleles with probability 0.25, one of the individual’s alleles chosen at random with probability 0.5, and none of the individual’s alleles with probability 0.25. We then chose the remaining allele(s) according to the weighted estimated allele frequency distribution given by Equation 1. We treated loci as independent, and we also treated alleles within loci as independent.

Our approach of simulating identity by descent between siblings according to the relatedness coefficients 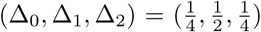 follows Rohlfs *et al*. (2012) in assuming no background identity by descent in the general population, or in other words, a coancestry coefficient *θ* = 0. However, unlike in Rohlfs *et al*. (2012), because the allele frequency distribution in our simulation was distinctive to each individual, it is possible that the method of simulation induces a level of coancestry *θ* > 0 between siblings of different sampled individuals comparable to that seen among individuals in the initial worldwide data set.

### Likelihood Ratios

#### Definition

We calculated likelihood ratios (LRs) for relationship hypotheses for each pair consisting of an individual and a simulated sibling. We performed this computation within each of the four prior population groups, following the procedure of Rohlfs *et al*. (2012). This step considered 94 × 940 pairs in A, 532 × 5320 in EMC, 269 × 2690 in EAO, and 83 × 830 in NA. We calculated the likelihood ratio

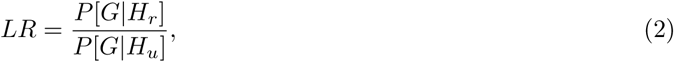

where *G* represents the multilocus genotype data for the pair, *H*_*r*_ is the hypothesis that the two individuals in the pair are related, and *H*_*u*_ is the hypothesis that they are unrelated. If we assume that all 13 CODIS loci are independent, then we can express Equation 2 as:

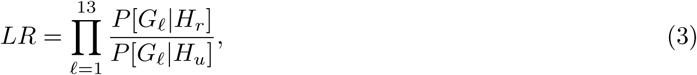

where *G*_*ℓ*_ represents the data at locus *ℓ, ℓ* = 1, …, 13. Evaluating Equation 3 entails inserting the coefficients of relatedness, which for siblings are 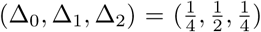 and for unrelated pairs are (Δ_0_,Δ_1_,Δ_2_) = (1,0,0):

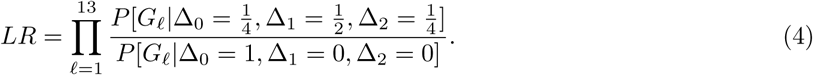

#### Calculation

Expressions for probabilities *P* [*G*_*ℓ*_|Δ_0_,Δ_1_,Δ_2_] depend on the combinations of alleles observed for pairs of individuals, on the allele frequencies assumed, and also on the assumed value of the coancestry coefficient *θ*, These expressions, originally derived by Fung *et al*. (2003), appear in Rohlfs *et al*. (2012), supplementary text, page 1 (in the last case, *P* (*A*_1_*A*_2_, *A*_3_*A*_4_ |Δ_2_,Δ_1_,Δ_0_), the equation is missing a coefficient of 4 that does not affect likelihood ratio computation). Following Rohlfs *et al*. (2012), we considered two values for the coancestry coefficient, *θ* = 0 and *θ* = 0.01. We include *θ* = 0.01 in the main text and *θ* = 0 in the supplement.

In evaluating the likelihoods, we considered a variety of ways of setting the allele frequencies (see Results).

#### Comparing Likelihood Ratio Distributions

To evaluate the difference between the likelihood-ratio distributions for true related and true unrelated individuals, we calculated the distinguishability measure 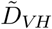 (Visscher and Hill, 2009; Rohlfs *et al*., 2012),

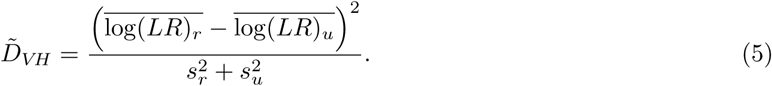

Here, 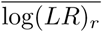 and log 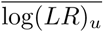 are sample means of the LR distributions for the true relatives and true unrelated pairs, respectively; 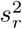 and 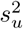 are the sample variances of the distributions of LRs for the true relatives and true unrelated pairs, respectively. A higher 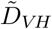 indicates that likelihood ratio distributions for true relatives and true unrelated individuals are more easily distinguished. We used base *e* for the logarithms in comparing 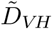 and in all other computations requiring logarithms.

The numbers of true relatives and true unrelated pairs in our simulation vary by assumed population group. Population A has 94 × 10 related pairs and 94 × (940 − 10) unrelated pairs. Population EMC has 532 × 10 related pairs and 532 × (5320 − 10) unrelated pairs. Population EAO has 269 × 10 related pairs and 269 × (2690 − 10) unrelated pairs. Population NA has 83 × 10 related pairs and 83 × (830 − 10) unrelated pairs.

### Gene Diversity

To assess a measure of the extent to which alleles in a population distinguish different individuals, we calculated the gene diversity, or expected heterozygosity, of each of the four populations. For each locus, the gene diversity is 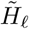 (Nei, 1987):

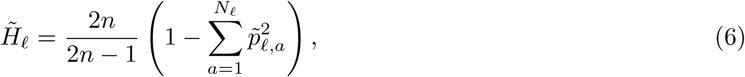

where *N*_*ℓ*_ is the number of distinct alleles at locus *ℓ*. Here, 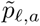 is the observed allele frequency of allele *a* at locus *ℓ* in the population and *n* is the sample size in the population for the locus. For each population, we averaged the observed gene diversity across all 13 CODIS loci to obtain 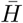. Note that no data were missing for the CODIS loci, so that a shared sample size *n* was used for all loci within each population.

### Coancestry Coefficients

We evaluated the degree of difference between pairs of populations in their allele frequency distributions. For this computation, we estimated *θ* for each pair of populations using the program GDA (Lewis and Zaykin, 2002). The calculation uses the estimator of Reynolds *et al*. (1983), as in Weir (1996), Equation 5.3. We present 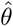 estimated using only the 13 CODIS loci as well as using all 791 loci.

### Data Availability

See Edge *et al*. (2017) for the data used in this study.

## Results

### Allele Frequencies from Predefined Populations

Following Rohlfs *et al*. (2012), we evaluated how misspecification of the assumed major population affects our ability to distinguish relatives from unrelated individuals. For each of our population groups, A, EMC, EAO, and NA, we computed likelihood ratios (LRs) for pairs of individuals and potential relatives using each of the four major populations’ estimated allele frequencies. We term this approach the *Predefined-Population* method of choosing the allele frequencies. When the assumed population matches the pair’s true population membership, we expect to more easily distinguish between true siblings and unrelated pairs compared to the cases in which the populations do not match.

In each panel of Figure 1, we show the distributions of LR values for true siblings and for true unrelated individuals, for a specific pair of true and assumed population memberships. For example, in the bottom leftmost panel, individuals are from the African population (A), and Native American allele frequencies (NA) are used to evaluate the likelihood ratios. The light green distribution is the density of log likelihood ratio values for true unrelated pairs, whereas the dark green distribution is for the true siblings. The black horizontal bars show the central 95% of each distribution. The plot uses a coancestry assumption of *θ* = 0.01. Plots along the diagonal of Figure 1 display the LR distributions for true siblings and unrelated individuals when the allele frequency assumption matches the true population. The off-diagonal plots show LR distributions for incorrect pairings of populations and allele frequency assumptions.

**Figure 1:**
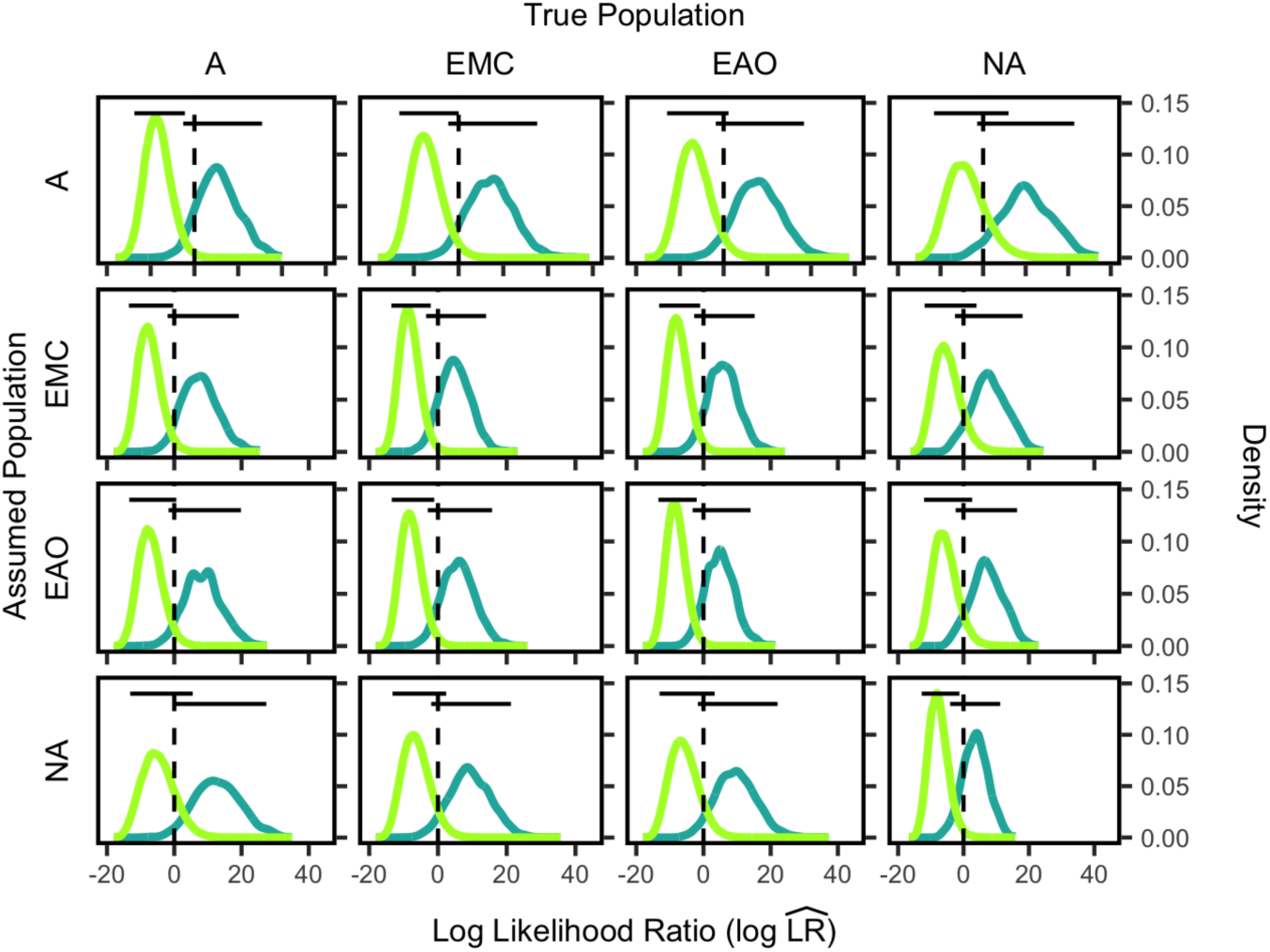
Log likelihood ratio 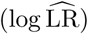 distributions for siblings and unrelated individuals by population group for allele frequencies chosen by the *Predefined-Population* method. Each plot shows the 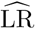 distributions for unrelated individuals in light green and true siblings in dark green, with each 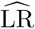 calculated from Equation 4. The dashed vertical lines indicate 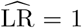 The horizontal lines show the central 95% of each distribution. Each distribution in the A column consists of 94 × (940 − 10) points and 94 × 10 points for the unrelated pairs and related pairs, respectively. Each distribution in the EMC column consists of 532 × (5320 − 10) and 532 × 10 pairs, respectively. Each distribution in the EAO column consists of 269 × (2690 − 10) and 269 × 10 pairs, respectively. Each distribution in the NA column consists of 83 × (830 − 10) and 83 × 10 pairs, respectively. A, African; EMC, European, Middle Eastern, and Central/South Asian; EAO, East Asian and Oceanian; NA, Native American.

Distinguishability between true relatives and unrelated individuals is higher when the matching allele frequencies are used rather than nonmatching allele frequencies, as shown by the minimal overlap between distributions in plots on the diagonal. In contrast, the off-diagonal plots have more overlap between the true-sibling and true-unrelated distributions. The specific distinguishability 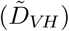 values are listed in Table 1, and they are consistent with an analogous analysis in Rohlfs *et al*. (2012), which also showed that distinguishability is highest when the assumed population matches the true population. Additionally, Rohlfs *et al*. (2012) found that distinguishability was lowest when Navajo was the true population, likely due to the relatively low genetic diversity within this population (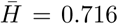 for the CODIS loci). Similarly, we also found the lowest distinguishability between pairs belonging to the Native American population. Rohlfs *et al*. (2012) found the highest distinguishability among African American samples, and we found the highest distinguishability among the Sub-Saharan African individuals, which have the highest diversity of all the populations we studied (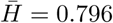 for the CODIS loci).

**Table 1:** 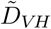 of major population groups, assuming allele frequencies from each major population group for the *Predefined-Population* method. 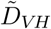 values are calculated using Equation 5 from the distributions plotted in Figure 1. A, African; EMC, European, Middle Eastern, and Central/South Asian; EAO, East Asian and Oceanian; NA, Native American.

|  | True Population |  |  |  |
| --- | --- | --- | --- | --- |
| Assumed Population | A | EMC | EAO | NA |
| A | 6.52 | 5.50 | 5.27 | 3.66 |
| EMC | 5.78 | 6.13 | 5.87 | 3.75 |
| EAO | 5.61 | 5.69 | 6.17 | 4.22 |
| NA | 4.55 | 4.72 | 4.39 | 5.28 |

### Allele Frequencies from Ancestry Inference

In the *Predefined-Population* method in Figure 1, specifying the correct-population allele frequencies clearly results in greater distinguishability than using misspecified allele frequencies. We hypothesized that further refining the allele frequencies using ancestry inference would also lead to higher distinguishability between related and unrelated individuals than using misspecified allele frequencies. Our *Ancestry-Estimation* method incorporates ancestry inference on query samples to create weighted allele frequency distributions for calculating LRs.

The most accurate ancestry estimates utilize all of the available data. Hence, we first performed STRUCTURE analysis of the 978 sampled individuals using all 791 STR loci. These “full-data” estimates are shown in Figure 2A. The clusters generally align with the four assumed populations, although each individual shows some amount of mixed cluster membership.

**Figure 2:**
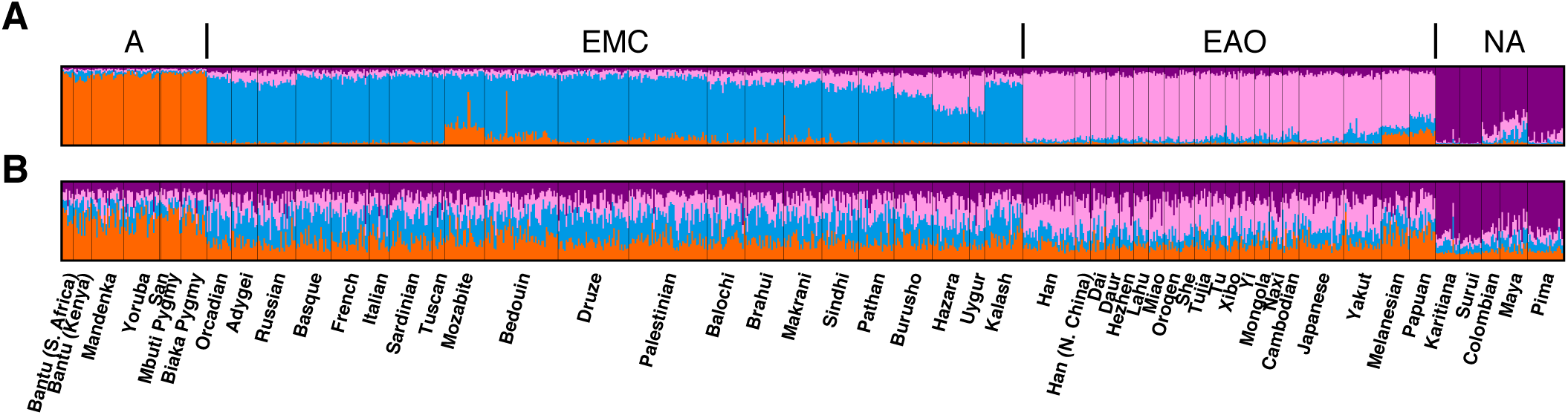
STRUCTURE-based inference with *K* = 4 clusters. (A) “Full-data” STRUCTURE results using all 791 loci. (B) “CODIS” STRUCUTRE results using only the 13 CODIS loci. Each color represents a different inferred cluster, and each cluster is generally associated with a prior population (orange, A; blue, EMC; pink, EAO; purple, NA).

However, in testing a query sample in a forensic context, ancestry would be estimated from fewer markers. Thus, we also performed STRUCTURE analysis using the 13 CODIS loci, as shown in Figure 2B. When we use the 13 CODIS loci instead of all 791 loci, each individual’s population membership is less clear, although each individual’s largest membership component generally matches that of the full-data STRUCTURE run. The analysis in Figure 2 is identical to that in Algee-Hewitt *et al*. (2016), except that one duplicated locus in Algee-Hewitt *et al*. (2016) was not duplicated in our analysis.

#### Likelihood Ratio Distribution

Next, for our *Ancestry-Estimation* method, we calculated weighted allele frequency distributions appropriate for each individual, weighting the inferred cluster allele frequencies by each individual’s inferred membership proportions, as in Equation 1. We then calculated likelihood ratios for each potentially related pair, as in Equation 4.

We tested three distinct scenarios for evaluating Equation 4. The first, “Full/Full,” uses the inferred allele frequencies and membership proportions from the full-data STRUCTURE results using all 791 loci in Figure 2A. This scenario is equivalent to possessing genotype data at all loci for both the query sample and global reference sample. The second, “Full/CODIS,” uses the inferred allele frequencies from the “full-data” STRUCTURE results, but uses the membership proportions from the “CODIS” STRUCTURE run using only the 13 CODIS loci in Figure 2B. This scenario amounts to having genotype data available at many loci for a global set of reference populations, enabling accurate inference of CODIS allele frequencies within inferred clusters. However, data are limited to the 13 CODIS loci for a query sample, so that ancestry estimates rely only on the CODIS loci. The third scenario, “CODIS/CODIS,” uses both the inferred allele frequencies and ancestry proportions from the “CODIS” STRUCTURE run. This scenario amounts to having data only at the 13 CODIS loci for both a global reference sample and for the query sample.

In each panel of Figure 3, we show the distribution of log likelihood ratios for true siblings in dark green and true unrelated individuals in light green, for a specific true population and a specific one of the three scenarios. The black horizontal bars show the central 95% of each distribution. For example, the top leftmost plot shows the density of LRs for true siblings and unrelated individuals in the African population (A), assuming the Full/Full scenario. The top row of Figure 3 shows the results for each population assuming the Full/Full scenario, the middle row shows the Full/CODIS scenario, and the bottom row shows the CODIS/CODIS scenario.

**Figure 3:**
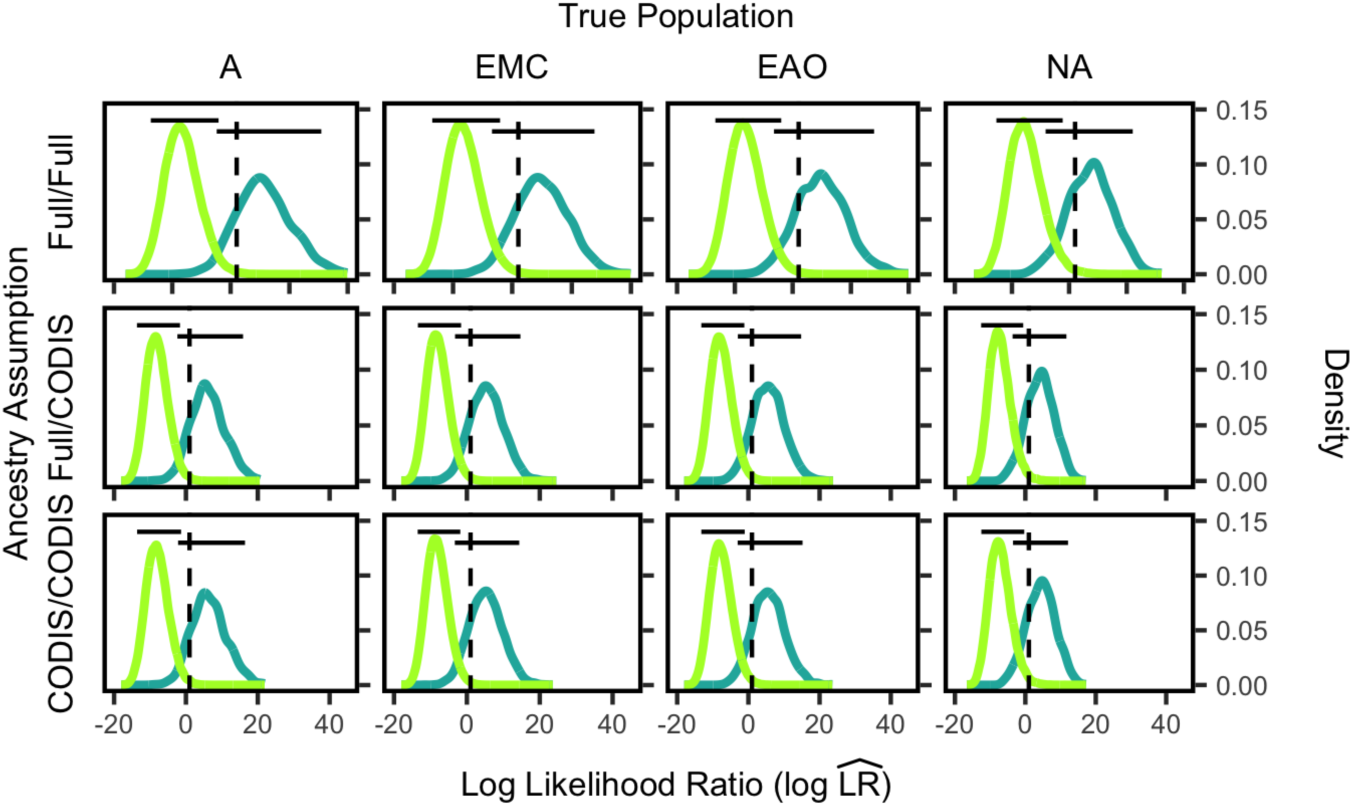
Log likelihood ratio 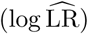 distributions for siblings and unrelated individuals by population group for allele frequencies chosen by the *Ancestry-Estimation* method. The labels on the left side indicate the scenario assumed, either Full/Full, Full/CODIS, or CODIS/CODIS. The figure design otherwise follows Figure 1.

The Full/Full assumption of Figure 3 produces the highest distinguishability between true siblings and unrelated individuals, as shown by the minimal overlap between the light green and dark green distributions. The CODIS/CODIS assumption generates the lowest distinguishability, as shown by the slightly higher overlap between the light green and dark green distributions. In other words, possessing as much data as possible (Full/Full) corresponds to a greater ability to distinguish true siblings and unrelated individuals. In contrast, the more limited data (CODIS/CODIS) is less successful in distinguishing true siblings and unrelated individuals.

#### Distinguishability

We next compared distinguishability assuming the *Ancestry-Estimation* method with distinguishability assuming the *Predefined-Population* method. Distinguishability values were calculated according to Equation 5 from the empirical distributions shown in Figures 1 and 3.

For the *Predefined-Population* method, for each of the true populations, we sort the values in Table 1 to rank the four ways of choosing the allele frequencies in decreasing order of distinguishability. The first of these four approaches, “Best-Specified Population,” uses an assumed population matching the individuals’ true population. There are then three misspecification scenarios; the identities of the assumed populations that correspond to each of these misspecification scenarios differ according to which true population is considered. Empirically, EMC is the second-best-specified population when the true population is A, but EAO is the second-best-specified population when the true population is NA, as shown in Table 1.

The results for each of the four *Predefined-Population* and three *Ancestry-Estimation* scenarios, ranked by highest average 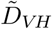 to lowest average 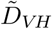 across rows of the table, appear in Table 2. The best-specified-population allele frequencies estimated from within a major population perform comparably to the Full/Full scenario, as they have similar 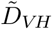 values across the row. The Full/CODIS assumption is the next highest, followed by the CODIS/CODIS assumption, which has a clearly lower average 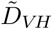 across the row than Full/CODIS. The three misspecified population assumptions all have much lower 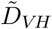 values across the row. Hence, misspecifying population allele frequencies generates a reduced ability to distinguish true relatives from unrelated individuals, in agreement with results from Rohlfs *et al*. (2012). Ancestry estimation to improve allele frequency estimates increases distinguishability over assuming an incorrect major population when the query individual’s major population membership is unknown.

**Table 2:** 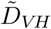for both methods, *Predefined-Population* and *Ancestry-Estimation*. **Full/Full**: Full-data allele frequencies and full-data ancestry proportions from STRUCTURE runs with 791 loci. **Full/CODIS**: Full-data allele frequencies from STRUCTURE runs with 791 loci and CODIS ancestry proportions from STRUCTURE runs with 13 CODIS loci. **CODIS/CODIS**: CODIS allele frequencies and CODIS ancestry proportions from STRUCTURE runs with 13 loci. **Best-Specified**: Allele frequencies from the assumed population to which the individuals and siblings belong. **Second-Best-Specified**: The second-highest distinguishability value from each column of Table 1, assuming the allele frequencies from the second-best assumed population. **Third-Best-Specified**: The third-highest distinguishability value from each column of Table 1, assuming the allele frequencies from the third-best assumed population. **Fourth-Best-Specified**: The lowest distinguishability value from each column of Table 1, assuming the allele frequencies from the fourth-best assumed population. 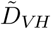 values are calculated using Equation 5 and the distributions plotted in Figures 1 and 3.

| Assumption | True Population |  |  |  |
| --- | --- | --- | --- | --- |
|  | A | EMC | EAO | NA |
| Best-Specified Population | 6.52 | 6.13 | 6.17 | 5.28 |
| Full/Full | 6.54 | 6.12 | 6.18 | 5.26 |
| Full/CODIS | 6.44 | 5.95 | 5.90 | 5.16 |
| CODIS/CODIS | 6.31 | 5.99 | 5.81 | 5.02 |
| Second-Best-Specified Population | 5.78 | 5.69 | 5.87 | 4.22 |
| Third-Best-Specified Population | 5.61 | 5.50 | 5.27 | 3.75 |
| Fourth-Best-Specified Population | 4.55 | 4.72 | 4.39 | 3.66 |

#### False Positive Rate and Power

Next, using the results in Figures 1 and 3, we assessed the false positive rate and power to distinguish relatives from unrelated individuals for both the *Ancestry-Estimation* and *Predefined-Population* methods. The left panel of Figure 4 shows the true positive rate for sibling detection as a function of the false positive rate, for pairs of individuals in the African population (A). Each color in this receiver-operating-characteristic (ROC) curve represents a different *Predefined-Population* or *Ancestry-Estimation* scenario. In these plots, curves that reach higher into the top-left corner of the plot have higher true positive rates of sibling detection at lower false positive rates. Each panel of Figure 4 shows results for a specified true population.

**Figure 4:**
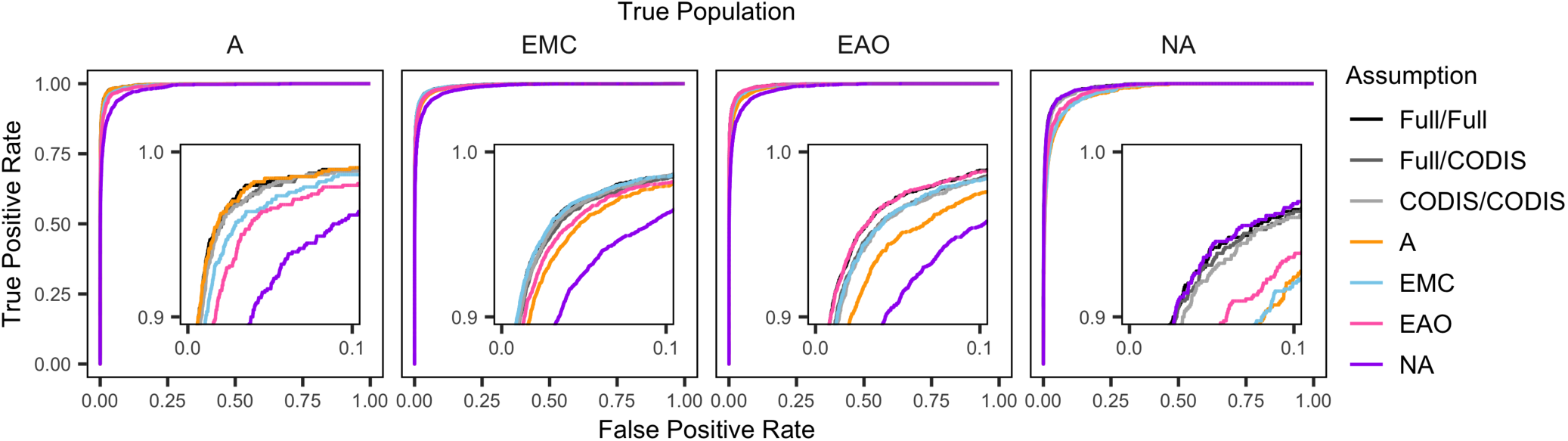
Receiver-operating-characteristic (ROC) curves showing true positive rate as a function of false positive rate in assigning individuals as siblings. The plots are calculated from the distributions in Figures 1 and 3. Each curve for A uses 94 × 940 pairs, each curve for EMC uses 532 × 5320 pairs, each curve for EAO uses 269 × 2690 pairs, and each curve for NA uses 83 × 830 pairs. The inset panels show the detail at the upper left corner of each plot.

The correct-population, Full/Full, Full/CODIS, and CODIS/CODIS assumptions largely overlap in this plot, irrespective of the true population. These assumptions have the highest area under the curve and are best able to distinguish true relatives from unrelated individuals. The misspecified-population scenarios, with lower distinguishability values, result in lower area under the curve.

### Gene Diversity

We expect to be able to distinguish relatives from unrelated individuals more easily when the corresponding allele frequency distribution has high rather than low variability. With low genetic diversity, individuals are more likely to have identical genotypes at a locus even when they are not close relatives.

Figure 5 shows distinguishability, 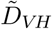 as a function of the average gene diversity across loci. The 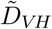 values are from Table 1, and the gene diversity is calculated according to Equation 6. The first three panels show the results for the three *Ancestry-Estimation* scenarios, and the last panel shows the results for the Best-Specified-Population scenario from the *Predefined-Population* method.

**Figure 5:**
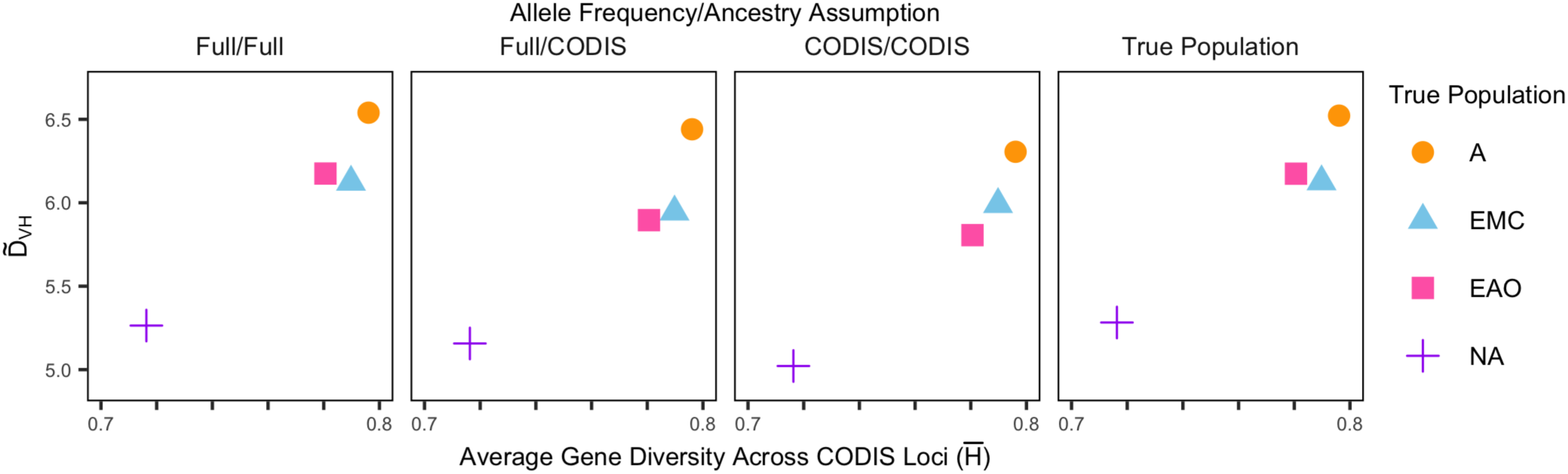
The empirical distinguishability 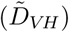 for siblings and unrelated individuals as a function of average gene diversity across the 13 CODIS loci, 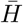. Points are colored according to the true population group. Each panel considers a different pair of assumptions about allele frequencies and ancestry in computing the likelihood ratios, as shown in Figures 1 and 3. 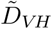 is computed from Equation 5 and 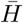 is computed from Equation 6.

Figure 5 shows that 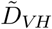 increases with gene diversity irrespective of the method used to evaluate LRs. The Native American (NA) population has the lowest gene diversity and 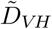, whereas the African population has the highest values of both quantities.

### Coancestry

We have shown that distinguishability is lower when the allele frequency assumption used to calculate likelihood ratios is incorrect. We quantify the degree of mismatch for misspecified and correctly specified allele frequency distributions using the coancestry coefficient, *θ*.

In Table 3, the upper triangle shows estimates of *θ* between populations using all 791 loci, and the lower triangle shows estimates of *θ* using the 13 CODIS loci. As a consequence of the high genetic diversity of CODIS loci informative for distinguishing individuals, the estimates using the 13 loci are smaller than the estimates using all 791 loci (Algee-Hewitt *et al*., 2016).

**Table 3:** 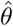 between population groups. The upper triangle was estimated using 791 loci, and the lower triangle was estimated using the 13 CODIS loci.

|  | A | EMC | EAO | NA |
| --- | --- | --- | --- | --- |
| A | - | 0.040 | 0.056 | 0.100 |
| EMC | 0.024 | - | 0.027 | 0.067 |
| EAO | 0.036 | 0.013 | - | 0.057 |
| NA | 0.081 | 0.058 | 0.053 | - |

Figure 6B shows 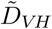 taken from Table 2, in relation to the estimated *θ*, taken from Table 3 under the scenarios from the *Predefined-Population* method. Figure 6A shows 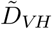 from Table 2 under the scenarios from the *Ancestry-Estimation* method, for comparison with the *Predefined-Population* case with correctly-specified populations (*θ* = 0) in Figure 6B. The left half of each circle is colored according to the prior population, and the right half is colored according to the assumed population or ancestry estimation scenario. Because 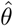 is calculated for each pair of populations, the two configurations of prior and assumed allele frequencies for a pair of populations lie at the same horizontal position in the plot.

**Figure 6:**
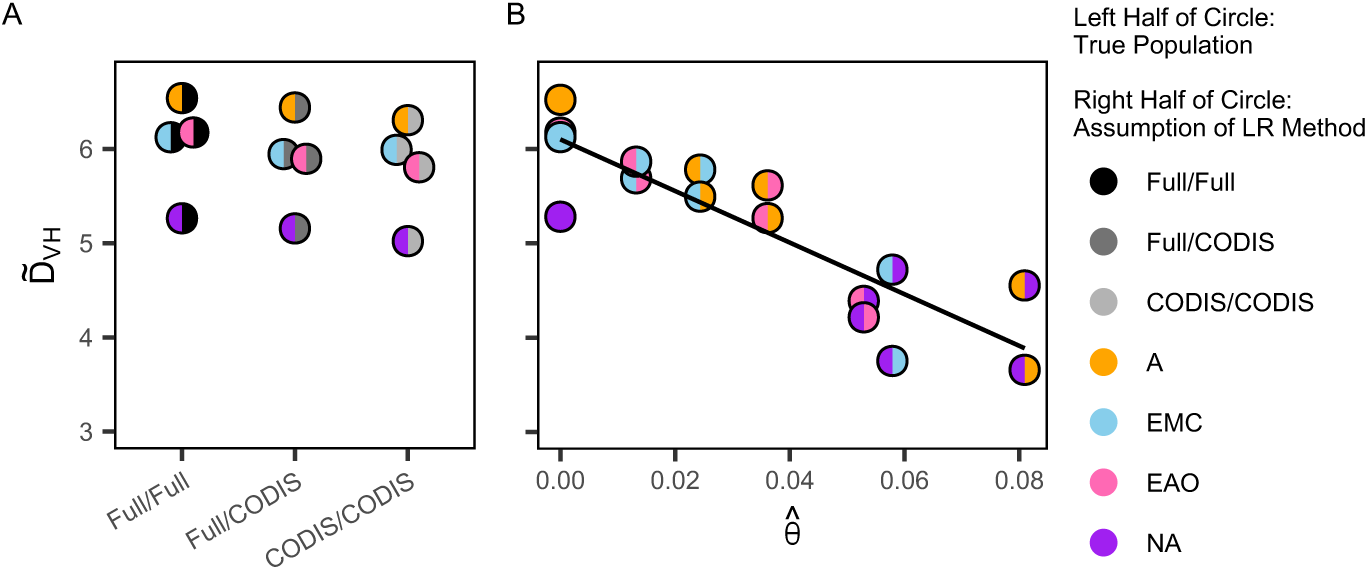
The empirical distinguishability 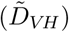 for siblings and unrelated individuals as a function of the estimated coancestry coefficient, 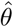, for pairs of populations, one reporting the true population and the other reporting the source for the allele frequencies. (A) Full/Full, Full/CODIS, and CODIS/CODIS *Ancestry-Estimation* scenarios. (B) *Predefined-Population* scenarios. The *θ* estimate is from the 13 CODIS loci only, as in the lower triangle of Table 3. The left color of each circle corresponds to the true population group, and the right color of each circle corresponds to the assumption used in the LR calculations. The Full/Full, Full/CODIS, and CODIS/CODIS cases are plotted separately in A for comparison with the *Predefined-Population* case with “correctly-specified” populations (single-color circles in B at 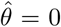). 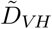 values are taken from Table 2. The equation of the regression line in B is 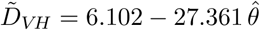.

As shown in Figure 6B, 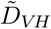 decreases with increasing 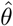 For the *Predefined-Population* method, the allele frequencies are increasingly misspecified as *θ* increases, decreasing our ability to distinguish true relatives from unrelated individuals.

## Discussion

In this study, we have analyzed methods for choosing allele frequencies for familial search in forensic genetics, comparing a new approach of using allele frequencies chosen from ancestry estimation in the query sample to use of allele frequencies from a predefined population. We have found that for the problem of distinguishing siblings from unrelated individuals, *Ancestry-Estimation* methods perform comparably to a *Predefined-Population* method that uses allele frequencies associated with the population of origin of the query sample (Table 2). The *Ancestry-Estimation* methods, however, avoid the high false positive rates that result from misspecifying the population of origin of the allele frequencies in the *Predefined-Population* method. In a forensic context, because genetic markers in query forensic profiles are always in principle available for ancestry estimation, the higher false positive rates resulting from the most extreme allele-frequency misspecifications can be avoided.

The study expands upon the work of Rohlfs *et al*. (2012), which characterized false positive rates in familial search using both allele frequencies matched by population to the query sample and misspecified allele frequencies. In a similar analysis using a different data set, we have replicated their results that false positive rates are substantially greater when the allele frequencies are misspecified (Figure 4), and that the increase in false positive rates increases with the degree of misspecification (Figure 6). Like Rohlfs *et al*. (2012), we found that distinguishability of relatives and unrelated individuals increases with gene diversity within populations, irrespective of the allele frequency scenario (Figure 5): as gene diversity increases across the four population groups, from Native Americans to Sub-Saharan Africans, the probability that a pair of non-siblings has a partial match decreases, increasing distinguishability.

Extending beyond the approach of Rohlfs *et al*. (2012) of considering allele frequencies from the population that matches the query sample and from each of several possible allele frequency misspecifications, we added three *Ancestry-Estimation* scenarios. All three scenarios produce greater distinguishability between siblings and unrelated individuals than use of misspecified allele frequencies, with values generally closer to those obtained for allele frequencies that match the query sample (Table 2). One of the *Ancestry-Estimation* scenarios, the CODIS/CODIS scenario, relies on allele frequencies and ancestry estimates obtained from the analysis of samples for which forensic markers have been gathered; this scenario is practical in principle in any case in which familial search is of interest and reference data are available on forensic genetic markers.

The Full/Full *Ancestry-Estimation* scenario, considering allele frequencies and ancestry estimates based on use of many more markers beyond the 13 forensic markers, produces distinguishability values that exceed those of the CODIS/CODIS scenario, and that are comparable to use of allele frequencies that match the query profile (Table 2). Interestingly, however, the Full/CODIS scenario, in which allele frequencies are estimated from STRUCTURE runs with a large number of markers but ancestry estimates are obtained from STRUCTURE runs only with the CODIS loci, has distinguishability more similar to the CODIS/CODIS case rather than to the Full/Full case, despite its use of STRUCTURE estimates of allele frequencies from a larger data set. It is possible that distinguishability does not increase because the allele frequency estimates and ancestry estimates rely on STRUCTURE runs that use different data, so that the estimated parameters are not taken from the same model.

We note several limitations. Because the analysis obtains allele frequencies based on individual multilocus genotypes rather than treating alleles as independent across loci, residual coancestry among the sampled individuals could affect our characterization of the parameter *θ*. Thus, although we simulated siblings using *θ* = 0, it is possible that the actual coancestry of pairs of unrelated individuals tested for relatedness exceeds 0. When we use *θ* = 0.01 to compute likelihood ratios, we obtain greater distinguishability between siblings and unrelated pairs than when we use *θ* = 0 (see Supplement). However, changing the choice of *θ* does not affect the relative position of the different allele frequency assumptions, so that our broad conclusions about the improvement of *Ancestry-Estimation* compared to allele frequency misspecification are unaffected.

We have only considered sibling relationships. In general, false positive rates are expected to be lower for parent-offspring relationships than with sibling relationships. Unlike for siblings, a parent and offspring share at least one identical allele at every locus; for an unrelated pair to achieve this level of sharing is more unlikely than to produce identity by chance at some of the loci, as in the case of tests for siblings or other relationships. Because our simulation approach, which did not take into account genotyping error, would find that nearly all unrelated pairs would be excluded as parent-offspring pairs, we focused on sibling relationships. However, our approach could potentially examine other relationships, such as half-siblings and first cousins.

An additional comment as that in the United States, for new samples starting in 2017, forensic profiles are generally obtained with 20 rather than 13 CODIS loci (Hares, 2015). With an increase from 13 to 20 loci, we expect that distinguishability will increase in all scenarios, including both *Predefined-Population* and *Ancestry-Estimation* methods. In particular, ancestry inference based on 20 loci will potentially improve, increasing distinguishability for the CODIS/CODIS scenario.

We selected *K* = 4 worldwide populations for choosing allele frequencies, based on the analysis of Algee-Hewitt *et al*. (2016), in which the CODIS loci enabled four clusters to be identified using STRUCTURE. The choice of the level of granularity for choosing allele frequencies in forensic problems requires careful consideration; we found here that for query samples, it is potentially valuable to consider allele frequencies as linear combinations of multiple potential source populations. Such an approach may be particularly valuable for recently admixed populations; as the populations in the study, from the Human Genome Diversity Panel, have not been selected for recent admixture, this hypothesis merits further investigation with alternative data sets.

The use of familial search methods in forensic genetics has generated much discussion. Expanding the search space from database entrants to their close relatives has the potential to identify the contributor of a query profile when no exact match to the profile is found (Bieber *et al*., 2006; Curran and Buckleton, 2008). However, use of familial search raises concerns about privacy, law, and policy related to such searches (Greely *et al*., 2006; Murphy, 2010); for example, the set of relatives accessible to such investigations might disproportionately represent disadvantaged populations to an unacceptable degree. A central parameter in such discussions is the false positive rate of familial search procedures, as the false positive rate affects the rate at which false-positive relatives of database entrants are subjected to intrusive investigations. Although our study suggests that an ancestry-inference procedure can potentially bound the false positive rate at values below those produced by the most serious misspecifications of allele frequencies, such reductions may continue to produce rates that are found to be intolerably high. In practical settings, it continues to be important to examine false positive rates for familial search procedures in relation to associated risks.

## Acknowledgements

We acknowledge NIH grant R01 HG005855 for support.

## Supplementary Information

**Figure S1:**
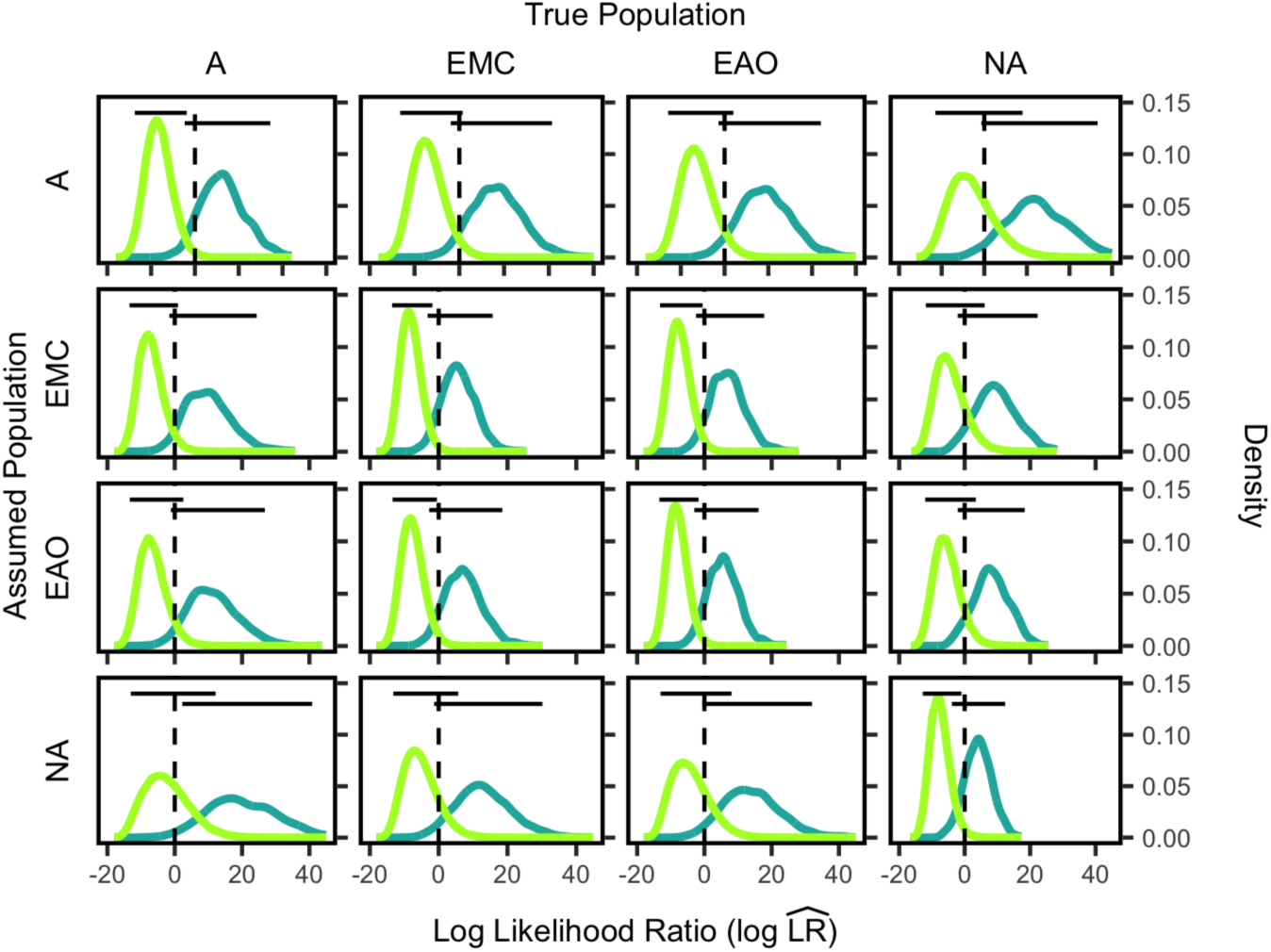
Log likelihood ratio 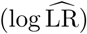 distributions for siblings and unrelated individuals by population group for allele frequencies chosen by the *Predefined-Population* method, with coancestry assumption *θ* = 0 (compare with Figure 1). Each plot shows the 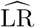 distributions for unrelated individuals in light green and true siblings in dark green, with each 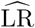 calculated from Equation 4. The dashed vertical lines indicate 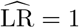 = 1. The horizontal lines show the central 95% of each distribution. Each distribution in the A column consists of 94 × (940 − 10) points and 94 × 10 points for the unrelated pairs and related pairs, respectively. Each distribution in the EMC column consists of 532 × (5320 − 10) and 532 × 10 pairs, respectively. Each distribution in the EAO column consists of 269 × (2690 − 10) and 269 × 10 pairs, respectively. Each distribution in the NA column consists of 83 × (830 − 10) and 83 × 10 pairs, respectively. A, African; EMC, European, Middle Eastern, and Central/South Asian; EAO, East Asian and Oceanian; NA, Native American.

**Figure S2:**
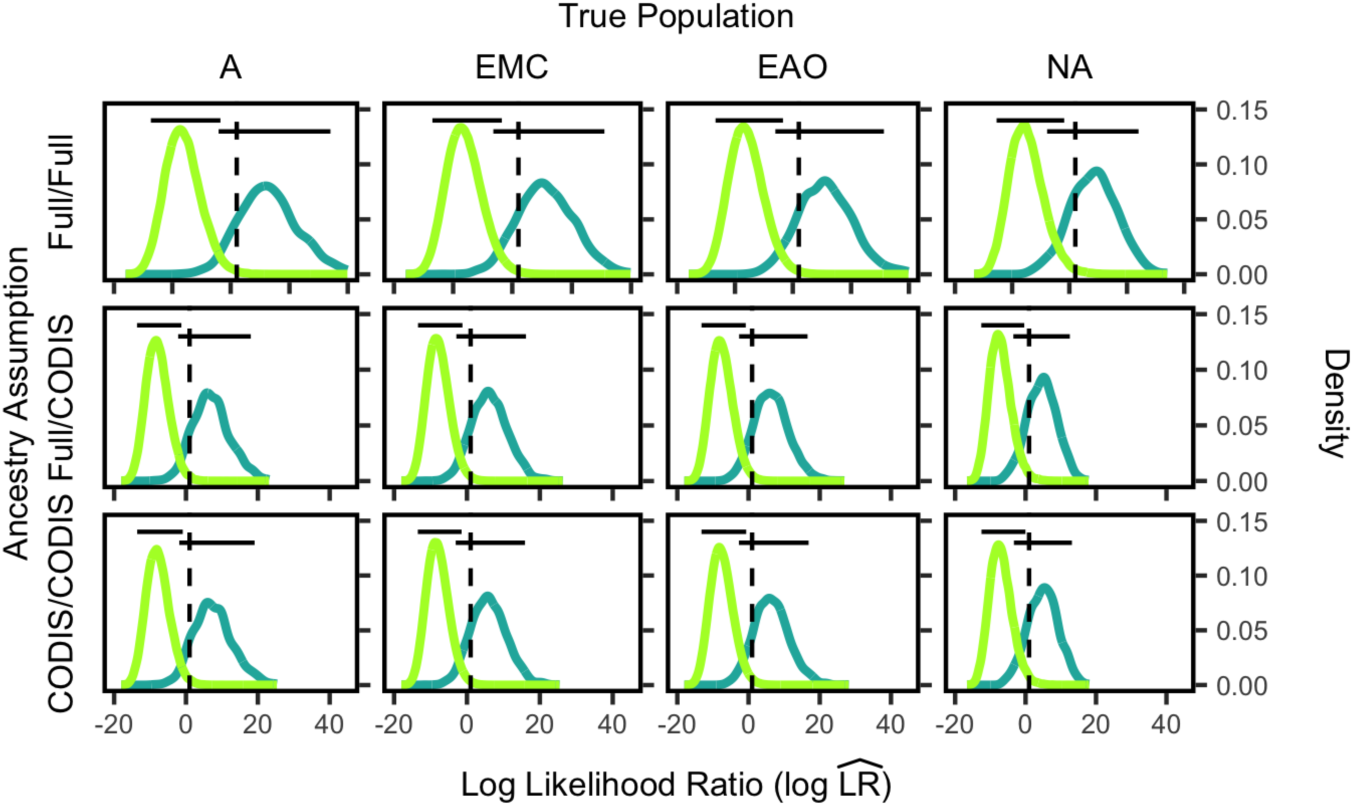
Log likelihood ratio 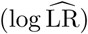 distributions for siblings and unrelated individuals by population group for allele frequencies chosen by the *Ancestry-Estimation* method, with coancestry assumption *θ* = 0 (compare with Figure 3). The labels on the left side indicate the scenario assumed, either Full/Full, Full/CODIS, or CODIS/CODIS. The figure design otherwise follows Figure S1.

**Figure S3:**
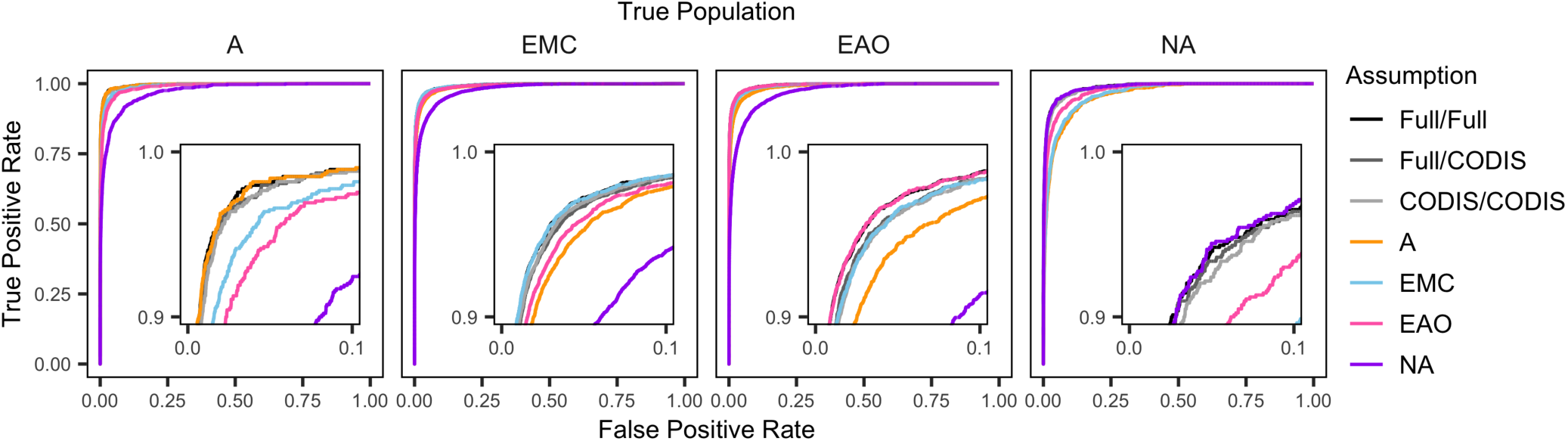
Receiver-operating-characteristic (ROC) curves showing true positive rate as a function of false positive rate in assigning individuals as siblings, with coancestry assumption *θ* = 0 (compare with Figure 4). The plots are calculated from the distributions in Figures S1 and S2. Each curve for A uses 94 × 940 pairs, each curve for EMC uses 532 × 5320 pairs, each curve for EAO uses 269 × 2690 pairs, and each curve for NA uses 83 × 830 pairs. The inset panels show the detail at the upper left corner of each plot.

**Figure S4:**
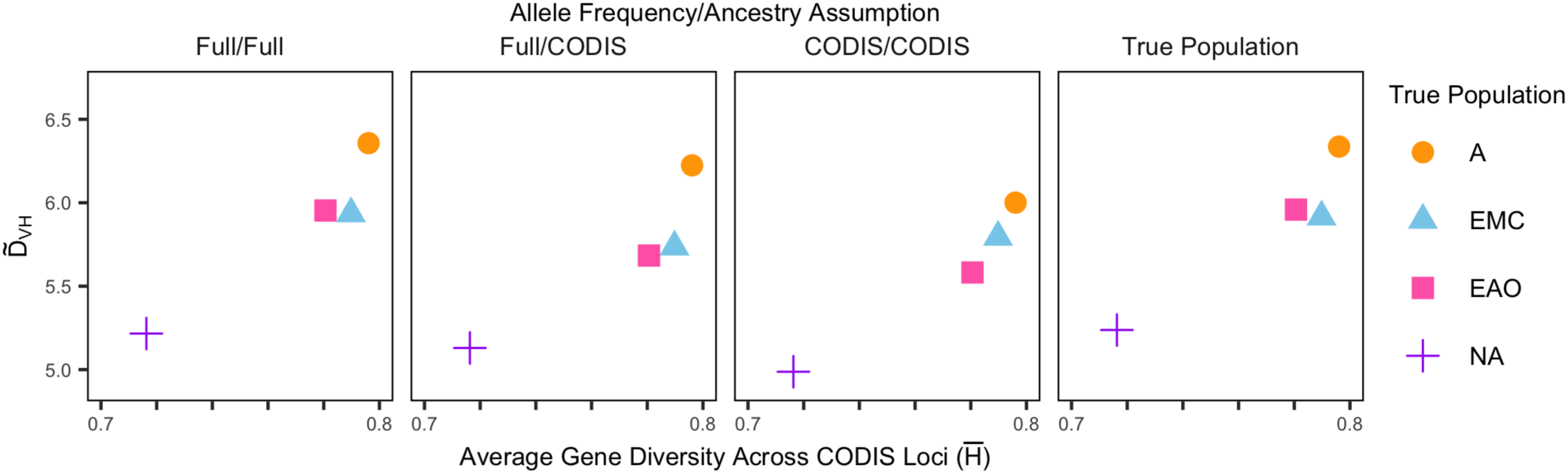
The empirical distinguishability 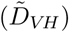 for siblings and unrelated individuals as a function of average gene diversity across the 13 CODIS loci, 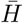, with coancestry assumption *θ* = 0 (compare with Figure 5). Points are colored according to the true population group. Each panel considers a different pair of assumptions about allele frequencies and ancestry in computing the likelihood ratios, as shown in Figures S1 and S2. 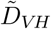 is computed from Equation 5 and 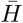 is computed from Equation 6.

**Figure S5:**
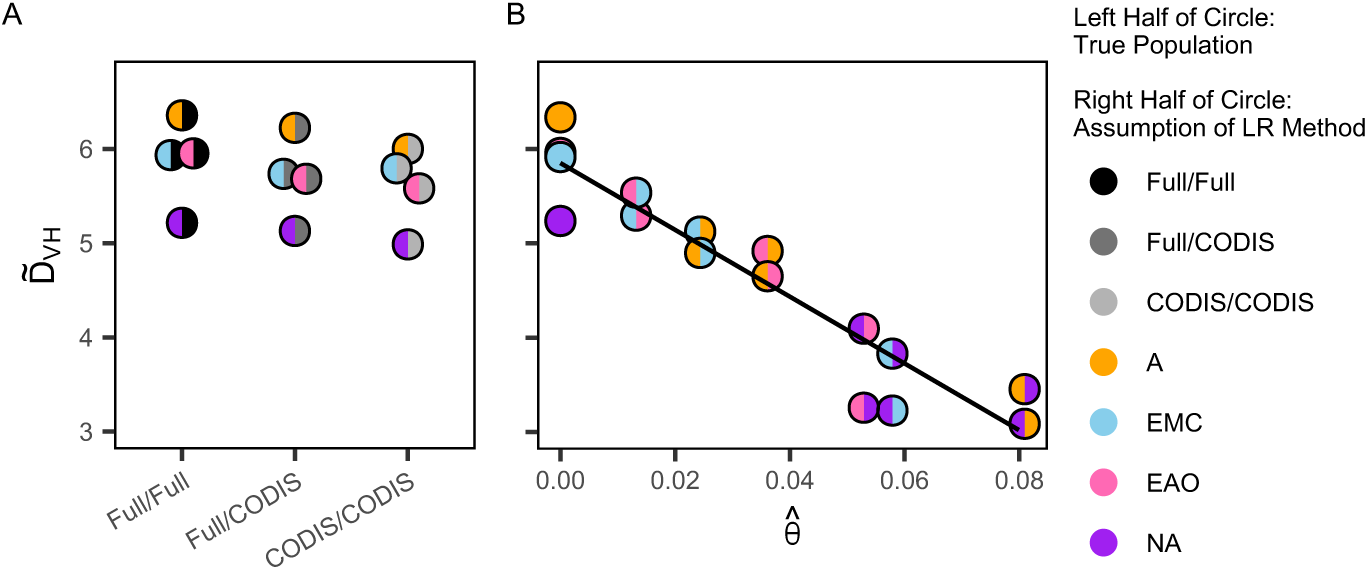
The empirical distinguishability 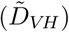 for siblings and unrelated individuals as a function of the estimated coancestry coefficient, 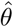, for pairs of populations, one reporting the true population and the other reporting the source for the allele frequencies. (A) Full/Full, Full/CODIS, and CODIS/CODIS *Ancestry-Estimation* scenarios. (B) *Predefined-Population* scenarios. The *θ* estimate is from the 13 CODIS loci only, as in the lower triangle of Table 3. The coancestry assumption is *θ* = 0 (compare with Figure 6). The left color of each circle corresponds to the true population group, and the right color of each circle corresponds to the assumption used in the LR calculations. The Full/Full, Full/CODIS, and CODIS/CODIS cases are plotted separately in A for comparison with the *Predefined-Population* case with “correctly-specified” populations (single-color circles in B at 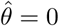). 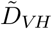 values are taken from Table S2. The equation of the regression line is 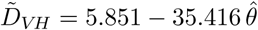.

**Table S1:** 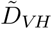 of major population groups, assuming allele frequencies from each major population group for the *Predefined-Population* method, with coancestry assumption *θ* = 0 (compare with Table 1). 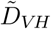 values are calculated using Equation 5 from the distributions plotted in Figure S1. A, African; EMC, European, Middle Eastern, and Central/South Asian; EAO, East Asian and Oceanian; NA, Native American.

|  | True Population |  |  |  |
| --- | --- | --- | --- | --- |
| Assumed Population | A | EMC | EAO | NA |
| A | 6.34 | 5.13 | 4.92 | 3.09 |
| EMC | 4.90 | 5.92 | 5.54 | 3.23 |
| EAO | 4.65 | 5.29 | 5.96 | 4.10 |
| NA | 3.45 | 3.83 | 3.26 | 5.24 |

**Table S2:** 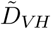 for both methods, *Predefined-Population* and *Ancestry-Estimation*, with coancestry assumption *θ* = 0 (compare with Table 2). **Full/Full**: Full-data allele frequencies and full-data ancestry proportions from STRUCTURE runs with 791 loci. **Full/CODIS**: Full-data allele frequencies from STRUCTURE runs with 791 loci and CODIS ancestry proportions from STRUCTURE runs with 13 CODIS loci. **CODIS/CODIS**: CODIS allele frequencies and CODIS ancestry proportions from STRUCTURE runs with 13 loci. **Best-Specified**: Allele-frequencies from the assumed population to which the individuals and siblings belong. **Second-Best-Specified**: The second-highest distinguishability value from each column of Table S1, assuming the allele frequencies from the second-best assumed population. **Third-Best-Specified**: The third-highest distinguishability value from each column of Table S1, assuming the allele frequencies from the third-best assumed population. **Fourth-Best-Specified**: The lowest distinguishability value from each column of Table S1, assuming the allele frequencies from the fourth-best assumed population. 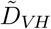 values are calculated using Equation 5 and the distributions plotted in Figures S1 and S2.

| Assumption | True Population |  |  |  |
| --- | --- | --- | --- | --- |
|  | A | EMC | EAO | NA |
| Best-Specified Population | 6.34 | 5.92 | 5.96 | 5.24 |
| Full/Full | 6.36 | 5.93 | 5.95 | 5.22 |
| Full/CODIS | 6.22 | 5.74 | 5.68 | 5.13 |
| CODIS/CODIS | 6.00 | 5.79 | 5.58 | 4.99 |
| Second-Best-Specified Population | 4.90 | 5.29 | 5.54 | 4.10 |
| Third-Best-Specified Population | 4.65 | 5.13 | 4.92 | 3.23 |
| Fourth-Best-Specified Population | 3.45 | 3.83 | 3.26 | 3.09 |

